# ATP-Citrate lyase fuels axonal transport across species

**DOI:** 10.1101/2020.07.07.192096

**Authors:** Aviel Even, Giovanni Morelli, Romain Le Bail, Michal Shilian, Silvia Turchetto, Loïc Broix, Alexander Brandis, Shani Inbar, Alain Chariot, Frédéric Saudou, Paula Dietrich, Ioannis Dragatsis, Bert Brone, Jean-Michel Rigo, Miguel Weil, Laurent Nguyen

**Author notes:** Corresponding authors: Laurent Nguyen, Miguel Weil. Equal contribution to the work (Aviel Even, Giovanni Morelli, Miguel Weil, Laurent Nguyen).

## Abstract

Microtubule (MT)-based transport is an evolutionary conserved processed finely tuned by posttranslational modifications. Among them, α-tubulin acetylation, which is catalyzed by the α-tubulin N-acetyltransferase 1, Atat1, promotes the recruitment and processivity of molecular motors along MT tracks. However, the mechanisms that controls Atat1 activity remains poorly understood. Here, we show that a pool of vesicular ATP-citrate lyase Acly acts as a rate limiting enzyme to modulate Atat1 activity by controlling availability of Acetyl-Coenzyme-A (Acetyl-CoA). In addition, we showed that Acly expression is reduced upon loss of Elongator activity, further connecting Elongator to Atat1 in the pathway regulating *α*-tubulin acetylation and MT-dependent transport in projection neurons, across species. Remarkably, comparable defects occur in fibroblasts from Familial Dysautonomia (FD) patients bearing an autosomal recessive mutation in the gene coding for the Elongator subunit ELP1. Our data may thus shine new light on the pathophysiological mechanisms underlying FD.

## Introduction

Axonal transport is an evolutionary conserved process that delivers cargoes to distant subcellular compartments. It is supported by molecular motors (kinesins and dyneins) that run along microtubule (MT) tracks and is particularly important for projection neurons that send axons to distant targets. MT-dependent transport contributes to neuronal development in growing dendrites and axons via slow axonal transport of cytoskeletal elements and, later, it sustains survival and homeostasis of neurons via fast axonal transport of organelles (mitochondria, lysosomes,..) and vesicles carrying various types of proteins (growth factors, synaptic proteins,..). Axonal transport defect is a hallmark of several neurodegenerative disorders, whose disruption affects neuronal function and survival (1-4). This is exemplified by loss of activity of the Elongator complex, which is associated with both neurodegeneration and axonal transport defects (5, 6). This molecular complex, conserved from yeast to human, is composed of two copies of six distinct protein subunits (Elp1 to Elp6) (7). Elp3 is the enzymatic core of Elongator and harbors two highly conserved subdomains, a tRNA acetyltransferase (8) and a S-adenosyl methionine binding domains (9). Elp1 is the scaffolding subunit of the complex but disruption of any of the Elongator subunits leads to comparable phenotype in eukaryotes, suggesting that all subunits are essential for the integrity and activity of the complex (10, 11). Elongator serves molecular functions in distinct subcellular compartments (12) but predominantly accumulates in the cytoplasm where it promotes the formation of 5-methoxycarbonylmethyl (mcm5) and 5-carbamoylmethyl (ncm^5^) on side-chains of the wobble uridines (U_34_) of selected tRNAs, thereby regulating protein translation (8, 13). Convergent observations support a role for Elongator in intracellular transport in the nervous system as Elp3 is enriched at the pre-synaptic side of neuromuscular junction buttons in flies, where its expression is required for synapse integrity and activity (14, 15). Elongator subunits are also detected in protein extracts from purified motile vesicles isolated from the mouse cerebral cortex (16), and they colocalize with the vesicular markers SV2 and RAB3A in human embryonic stem cells derived neurons (17). In humans, mutation of the gene coding for ELP1, underlies Familial dysautonomia (FD), a devastating disease that mostly affects the development and survival of neurons from the autonomic nervous systems (18, 19). Moreover, other neurological disorders are also associated with mutations affecting the expression or the activity of Elongator (5, 20-23). Experimental data on animal models show that interfering with Elongator activity promotes early developmental and progressive neurodegenerative defects, including poor axonal transport and maintenance (24-27). At the molecular level, loss of Elongator indeed correlates with poor acetylation of the α-tubulin lysine 40 (K40) in neuronal microtubules (MT) (6, 25, 28). This post-translational modification (PTM) modulates axonal transport by facilitating the recruitment of molecular motors to MTs (29) and the loading of motile vesicles on motors (6). The acetylation of MTs relies mostly on a vesicular pool of tubulin acetyltransferase 1 (Atat1) (30) that catalyzes the transfer of acetyl groups from Acetyl-Coenzyme-A (Acetyl-CoA) onto lysine 40 (K40) of α-tubulin (31, 32). How loss of Elongator affects MT acetylation, and whether a functional correlation between Elongator and Atat1 exists, remains to be explored.

Here, by combining cellular and molecular analyses in mouse cortical neurons *in vitro* and fly larva motoneurons *in vivo*, we show that loss of Elongator impairs axonal transport and acetylation of α-tubulin across species. Moreover, our findings support a common pathway where Elongator modulates Atat1 activity by regulating the expression of the ATP-citrate lyase (Acly), which produces acetyl-coA and thus provides acetyl groups for Atat1 activity towards MTs. Importantly, analysis of primary fibroblasts from FD patients show molecular defects comparable to those observed in mice and fly projection neurons depleted in Elongator. Therefore, our data may shine new light on the pathophysiological mechanisms underlying FD and other neurological disorders resulting from impaired Elongator activity.

## Results

### Elongator and Atat1 cooperate in a common pathway to regulate the acetylation of α-tubulin and axonal transport

In order to understand how Elongator controls tubulin acetylation and MT-based transport, we first performed complementary experiments in distinct animal models that lack Elongator activity. We compared the level of *α*-tubulin acetylation in cultured cortical projection neurons (PN) that were isolated from embryonic (E) day 14.5 WT or Elp3cKO mouse embryos (conditional loss of *Elp3* in cortical progenitors upon breeding *Elp3*lox/lox (33) and FoxG1:Cre (34) transgenic mice). The axon of *Elp3* cKO PNs displayed 50% reduction of acetylation of α-tubulin (Figure 1a). We next assessed axonal transport in PNs that were cultured in microfluidic devices for five days and incubated with specific dyes to track lysosomes (LysoTracker®) and mitochondria (MitoTracker®) movements by time-lapse videomicroscopy (Figure 1b). *Elp3* cKO PNs showed a significant reduction of average and moving velocities of lysosomes and mitochondria along axons, which correlated with an increase of their pausing times, as compared to WT PNs. Moreover, the moving velocities of lysosomes and mitochondria were reduced in both anterograde and retrograde directions (Figure S1a-S1g). These results confirmed that loss of Elongator activity leads to defects of α-tubulin acetylation and molecular transport in cortical neurons. Since a vesicular pool of Atat1 promotes the acetylation of α-tubulin, thereby axonal transport in cortical PNs (30), we postulated that Atat1 and Elongator might contribute to axonal transport via a shared molecular pathway. To test this hypothesis, we infected cortical PNs from E14.5 WT and *Atat1* KO mice with lentiviruses expressing either sh-Elp3 or sh-control (Figure S1h-i), and we cultured them in microfluidics devices for five days (Figure 1b). Targeting Elp3 in WT neurons impaired lysosomes and mitochondria transport across PN axons. However, we did not observe any additive effect on MT acetylation or axonal transport kinetics upon reduction of Elp3 expression in *Atat1* KO PNs (which show reduced acetylation of α-tubulin; Figure S1j) (Figure 1c-e, Figure S1k-n). Comparable observations were made *in vivo* in motoneurons (MNs) from 3^rd^ instar larvae of *Drosophila melanogaster* obtained by crossing UAS:*Elp1* or UAS:*Elp3 RNAi* fly lines with D42:Gal4 line specific for MNs (further called *Elp1* KD and *Elp3* KD). The knockdown efficiency of the UAS:*Elp1 and* UAS:*Elp3* in postmitotic neurons was validated in adult fly heads (Elav:Gal4 driver; Figures S1o-p). We first confirmed that the level of α-tubulin acetylation was reduced in *Elp1* and *Elp3* KD 3^rd^ instar larvae MNs, a defect genetically rescued by expressing human (h) ELP3 or by co-targeting the main α-tubulin deacetylase HDAC6 with RNAi (*Elp1;Hdac6* KD, *Elp3;Hdac6* KD) (Figure 1f). Moreover, knocking down both *Elp3* and *Atat1* (*Elp3;Atat1* KD) did not further reduce the acetylation levels of α-tubulin as compared to *Elp3* KD alone (Figure 1f; p=0.259 and p=0.775, respectively). *In vivo* time-lapse recordings of axonal transport of synaptotagmin-GFP (SYT1-GFP) in MNs from anesthetized fly 3^rd^ instar larvae (Figure 1g) correlated with levels of α-tubulin acetylation (Figure 1f). We observed a reduction of both average and moving velocities of SYT1-GFP vesicles together with an extension of their pausing time, both in *Elp1* KD or *Elp3* KD larvae, and with no cumulative defects in *Elp3;Atat1* KD flies (Figures 1h-j, Figure S1q). Moreover, targeting *Hdac6* fully rescued the observed axonal transport defects observed in Elp1/3 KD in 3^rd^ instar larvae MNs (*Elp1;Hdac6* KD, *Elp3;Hdac6* KD) (Figures 1h-i, Figure S1q).

**Figure 1.**
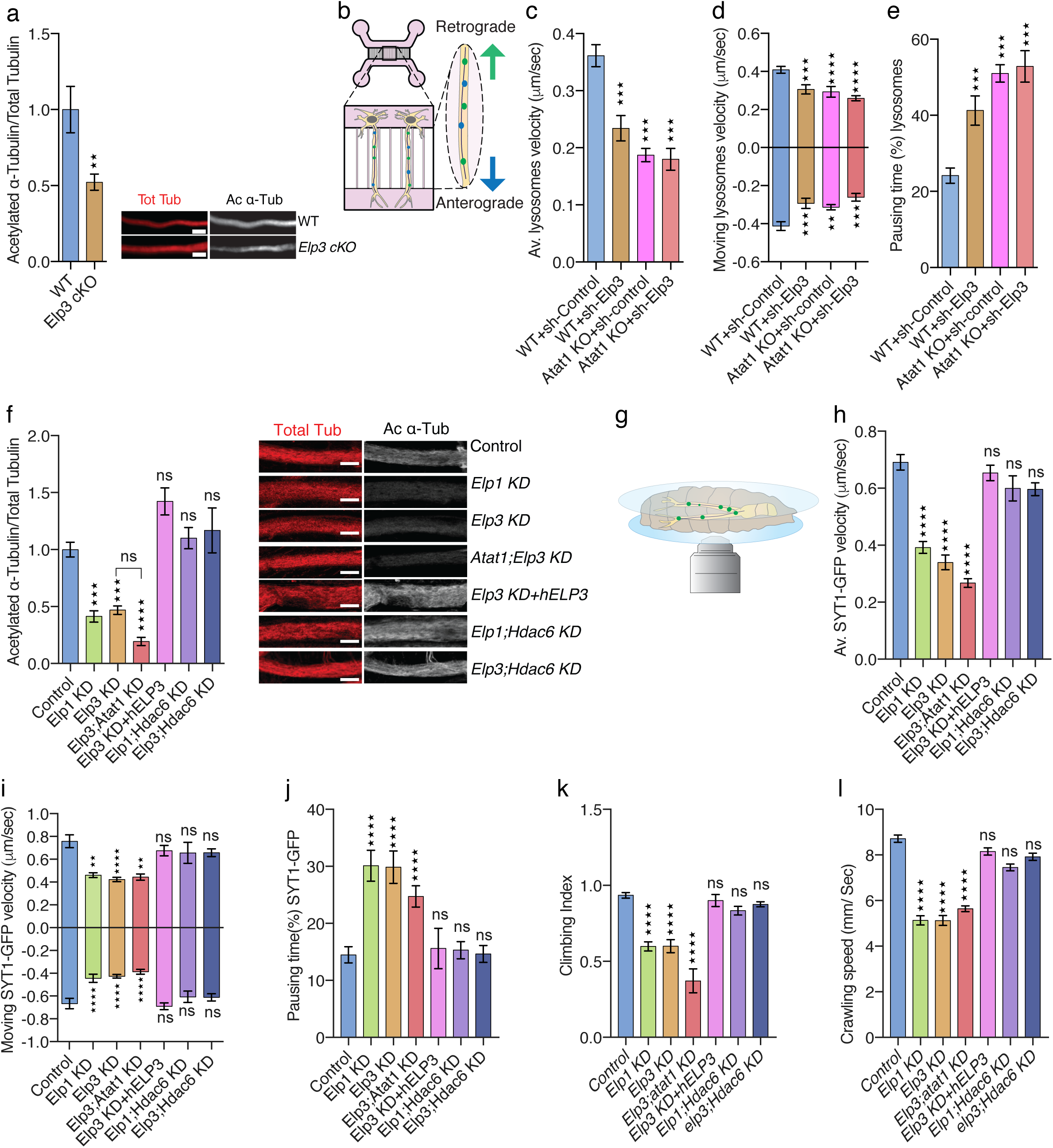
Reduction of Elongator activity leads to axonal transport defects in mouse and fly. (**a)** Immunolabelings and quantification of acetylated α-tubulin (Ac α-Tub) and total tubulin (Tot Tub) in axon of E14.5 WT and *Elp3 cKO* mice cortical neurons cultured for 5 DIV in microfluidic devices. Scale bar is 10 μm **(b**) Experimental set up to record axonal transport by time-lapse microscopy in E14.5 mice cortical neurons cultured for 5 DIV in microfluidic devices. (**c-e**) Histograms showing axonal transport of lysosomes (LysoTracker®) in WT or *ATAT1 KO* cultured neurons infected with sh-Control or sh-*Elp3* to analyze average (av.) velocity (**c**), moving velocity (**d**) and percentage of pausing time (**e**). (**f-l**) Study of *Drosophila melanogaster* expressing RNAi under a motoneuron-specific driver (D42:GAL4); control, *Elp1* KD, *Elp3* KD, *ATAT1; Elp3* KD, *Elp3* KD + human *Elp3, Elp1;Hdac6* KD and *Elp3;Hdac6* KD. (**f)** Immunolabelings and quantification of acetylated α-tubulin (Ac α-Tub) and total tubulin (Tot Tub) in motoneuron axon of 3^rd^ instar larvae, genotype as indicated. Scale bar is 10 μm. (**g**) Experimental set up for *in vivo* time-lapse recording of Synaptotagmin-GFP (Syt1-GFP) axonal transport in 3^rd^ instar larvae motoneurons, to analyze average (av.) velocity (**h**), moving velocity (**i**) and percentage of pausing time (**j**). (**k-l**) Locomotion assays: histograms of the climbing index of adult flies (**k**) and the crawling speed of 3^rd^ instar larvae (**l**). Description of graphical summaries here within are histograms of means ± SEM, while statistical analyses of: (**a**) is two-tailed t-test, (**c, d, e, f, h, i, j, l**) are Kruskal-Wallis test, and (**k**) is one-way analysis of variance (ANOVA). Specifically, [(**a**) p=0.0070, t=2.812, df=5; (**c**) p<0.0001 and K=80.47; (**d**) p<0.0001, K=34.30 and p<0.0001, K=23.46 for anterograde and retrograde, respectively, (**e**) p<0.0001 and K=73.34, (**f**) p<0.0001 and K=88.16, (**h**) p<0.0001 and K=223.6, (**i**) p<0.0001, K=110.3 and p<0.0001, K=45.69 for anterograde and retrograde, respectively, (**j**)p<0.0001, K=46.06, (**k**) p<0.0001, F=28.74; (**l**) p<0.0001, K=86.04. In addition, post hoc multiple comparisons for (**c, d, e, f, h, i**, **j**, **l**) Dunn’s test, for (**k**) Dunnet’s test and are **p < 0.01, ***p < 0.001, and ****p < 0.0001. The total number of samples analyzed were as follows: (**a**) 25-28 neurons from 3 different mice per group; (**c-e**) 58 to 180 vesicles coming from 3 different mice per group (at least 5 images per animal); (**f**) 14 to 32 motoneurons from at least 5 larvae per group; (**h-j**) 81-264 vesicles tracks from 7-12 larvae per group; (**k**) 6 to 15 test per group containing 10 adult flies; (**l**) 12 to 21 larvae per group.

Interfering with Elongator activity affects codon-biased translation that can ultimately lead to protein aggregation (35), thereby blocking axonal transport (36). However, we did not detect significant protein aggregation in the axon of either cultured cortical neurons from *Elp3* cKO mice (Figure S1r) or motoneurons from *Elp1* or *Elp3* KD 3^rd^ instar larvae in vivo (Figure S1s), while axonal aggregates formed upon blocking the proteasome activity with MG132 incubation. Altogether, our results suggest that the transport defects observed in the axon of Elongator deficient neurons unlikely result from a local accumulation of protein aggregates. Since impaired axonal transport in fly MNs leads to locomotion defects (1, 37), we measured larvae crawling speed and adult flies climbing index as functional readouts of axonal transport activity in MNs (30, 37). These parameters were affected upon depletion of either *Elp1* or *Elp3* (Figures 1k-l), and are likely the result of axonal transport defects, as we did not observe morphological changes at the synapses of the neuromuscular junctions (Figure S1u).

Elongator shares high amino acid similarity across species (Figure S1t). Expression of human *ELP3* in MNs of 3^rd^ instar larvae of *Elp3* KD flies (*Elp3* KD+*ELP3*) rescued α-tubulin acetylation (Figure 1f), axonal transport parameters (Figure 1h-j, Figure S1q) and locomotor behavior defects (Figures 1k-l), indicating that ELP3, and by extension Elongator, has a conserved evolutionary role from human to fly in axonal transport. Moreover, we show that loss of Elongator activity does not worsen the axonal phenotype of *Atat1* KO neurons, further suggesting that these molecules may cooperate in a common molecular pathway to modulate MT acetylation and axonal transport (31, 32).

### Loss of Elongator impairs the production of acetyl-CoA, thereby interfering with Atat1-mediated MT acetylation

Since *Hdac6* knockdown rescued the axonal transport observed in Elongator deficient neurons (Figure 1f-1k), we tested whether these defects may arise from a change of expression or activity of Hdac6 and/or Atat1, the enzymes that control α-tubulin de/acetylation, respectively (38, 39). Since no change in Atat1 and Hdac6 expression in cortical extracts from *Elp3* cKO and WT littermate newborn mice (Figures 2a-c) were observed, the activity of Atat1 was measured by using an *in vitro α*-tubulin acetylation assay (30). For this assay, pre-polymerized unacetylated MTs from HeLa cells were incubated with acetyl-CoA and P0 mouse brain extracts from *Elp3* cKO or WT littermate controls (Figures S2c-S2d) to assess the level of acetylation of α-tubulin. In strike contrast to brain extracts from *Atat1* KO mouse pups, we did not observe any differences in MT acetylation levels between cortical extracts from *Elp3* cKO and WT mice (Figure 2d). Moreover, the deacetylation activity of Hdac6 towards MTs, assessed *in vitro* by incubating free acetylated *α*-tubulin from bovine brains with cortical extracts from newborn *Elp3* cKO or WT littermate controls (38), was comparable between conditions (Figure 2e). Altogether, these results suggest that loss of Elongator activity does not impair the acetylation of α-tubulin by changing the expression or activity of Hdac6 or Atat1. Since the vesicular pool of Atat1 is predominant (30), we performed sub-fractionation of cortical lysates from adult mice lacking Elp1 (27), the scaffolding protein of the Elongator complex (*Elp1* cKO mice, Figure S2b) and analyzed their “vesiculome” using LC-MS/MS (Figure 2f). While Atat1 expression remained unchanged, the level of the ATP–citrate lyase (Acly), which converts glucose-derived citrate into acetyl-CoA (40) was significantly reduced upon loss of Elp1 expression (Figures 2g-2h). Acly generates acetyl-CoA and provides acetyl groups to Atat1 for acetylation of α-tubulin K40 in MTs (32). We thus hypothesized that the acetylation of α-tubulin could be regulated by homeostatic changes in Acly-dependent acetyl-CoA concentration, as previously observed for histone acetylation (41). Specifically, we postulated that loss of Elongator could interfere with MT acetylation and axonal transport via reduction of Acly expression. To test this hypothesis, we performed an *in-vitro* acetyl-CoA production assay (malate dehydrogenase coupled assay) (41), where subcellular fractions (T; total, S2: cytosolic + vesicular, S3: cytosolic, P3: vesicular; see figure 2f) of cortical brain extracts from *Elp3* cKO and WT littermate newborns was mixed with Acly substrates (ATP, CoA and citrate) (Figure 2i). The vesicular fraction (P3), which is strongly enriched into Elongator subunits (Elp1 and Elp3) Atat1 and Acly (Figures 2K, S2a), more efficiently produced acetyl-coA (Figures 2i-j). While cortical extracts from *Elp3* cKO E14.5 mouse embryos (Figures S2c-S2d) showed no change of *Acly* transcription (Figure 2l), we observed a reduction of Acly expression together with a correlative decrease of MTs acetylation (Figure S2d). Indeed, the analysis of the *Elp3* cKO and WT newborn littermates show a robust decrease of Acly protein level in both the cytosolic (*S3*, Figure 2m) and the vesicular fractions (*P3*, Figure 2n). Moreover, we detected a reduction of acetyl-CoA level in cortical extracts from *Elp3* cKO mice, as compared with WT newborn littermates by using LC-MS metabolic analysis (Figure 2o). Altogether, these results support an existing cooperation between Elongator and Atat1 in MT-acetylation via the regulated expression of the rate-limiting enzyme Acly, which provides acetyl-group donors to Atat1 for catalyzing *α*-tubulin acetylation in neurons.

**Figure 2.**
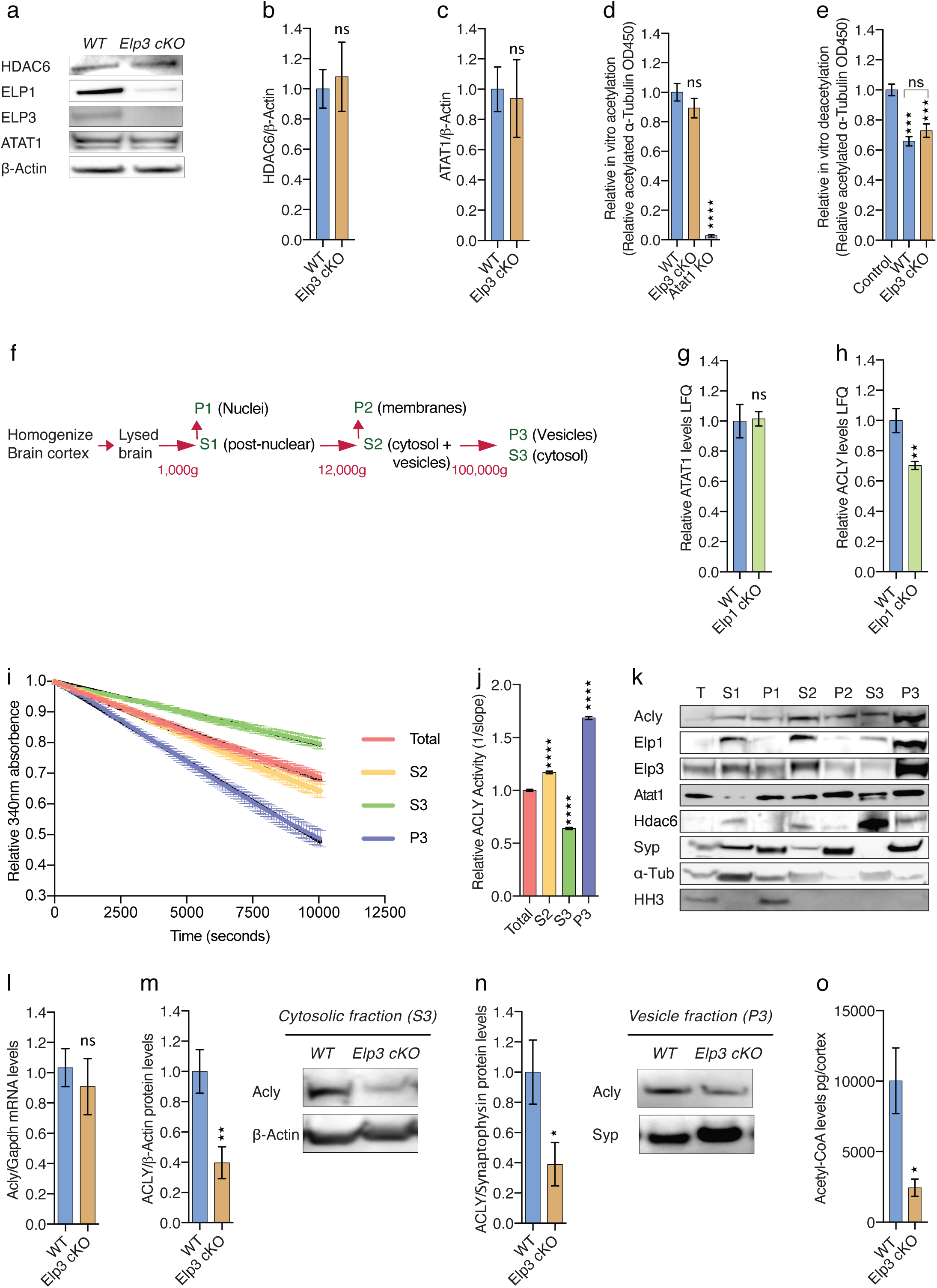
Elongator deletion impairs tubulin acetylation and axonal transport via reduction of Acly-dependent acetyl-coA production. (**a-c**) Immunoblotting to detect Atat1, Hdac6, Elp3 and ß-Actin cortical extracts from newborn WT and *Elp3 cKO* mice and histograms of proportion of Atat1(**b**) and Hdac6 (**c**) expression to ß-Actin. (**d**) In vitro acetylation assay of non-acetylated MTs from HeLa cells incubated for 2 hours with extracts of brain cortices isolated from WT, *Elp3 cKO* or *Atat1 KO* mice. (**e**) In vitro deacetylation assay of endogenously acetylated bovine brain tubulin incubated for 4 hours with extracts of cortices isolated from WT, *Elp3 cKO* or without tissue extract (control). (**f**) Experimental pipeline for subcellular fractionation of mouse cortical extracts: T, total; S1, postnuclear; P1, nuclear; S2, cytosol and vesicles; P2, larges membranes; S3, cytosol; and P3, vesicles. (**g-h**) Histogram of relative Atat1 and Acly label free quantification (LFQ) intensities analyzed by LC-MS/MS of P3 fraction from WT and *Elp1 cKO* mice. (**i-j**) Analysis of Acly activity by malate dehydrogenase coupled method performed in WT and *Elp3 cKO* brain cortex lysates. (**i**) Histogram of relative Acly activity over assay (i) and of slopes (j) from the linear phase of the reaction. (**k**) Subcellular fractionation (T, S1, P1, S2, P2, S3, P3) of mouse brain cortex immunoblotted with specific antibodies for Acly, Elp1, Elp3, Atat1, Hdac6, synaptophysin (Syp), α-tubulin (α-Tub),and Histone H3 (HH3). (**l**) qRT-PCR analysis of *Acly* mRNA in cortical brain extracts of *Elp3 cKO* or WT littermate mice. (**m-n**) Immunoblotting and quantification of cytosolic (**m**) and vesicular (**n**) fractions of Acly protein from newborn WT and *Elp3 cKO* mice brain cortices. (**o**) LC-MS quantification of Acetyl-CoA levels in WT and *Elp3 cKO* P0 mice brain cortex lysates. Description of graphical summaries here within are histograms of means ± SEM, while statistical analyses of (**b, c, m, n**) are two-tailed t-test, (**g, h, l, o**) are Mann-Whitney test, (**d, e, j**) are one-way analysis of variance (ANOVA), (i) is RM two-way ANOVA. Specifically, [(**b**) p= 0.7505, t=0.3248, df=13; (**c**) p= 0.8411, t=0.2093, df=6; (**d**) p<0.0001, F=85.29; (**e**) p<0.0001, F=21.30, (**g**) p=0.7619, U=10, (**h**) p=0.0087, U=2, (**i**) p<0.0001, F_Interaction_ (294, 882) = 68.07; (**j**) p<0.0001, F=1352; (**l**) p=0.5714, U=5; (**m**) p=0.0145, t=2.855,df=12; (**n**) p=0.0428, t=2.470, df=7; (**o**) p=0.0159, U=0. In addition, post hoc multiple comparisons for (**d**,**e**,**j**) are Holm-Sidak test, and are *p<0.05, **p < 0.01, ***p < 0.001, and ****p < 0.0001. The total number of samples analyzed were as follows: (**b**) 6-9 animals per group replicated 3 times; (**c**) 4 animals per group replicated 3 times; (**d**) 4-8 animals per group replicated 3 times; (**e**) 4-5 animals per group replicated 3 times; (**g-h**) 6 animals per group; (**i-j**) 4 animals per group subfraction preparation replicated 3 times for 99 time points (**s**); (**l**) 3-5 animals per group; (**m-n**) 5-9 animals per group; (**o**) 4-5 animals per group.

### Expression of Acly rescues both the level of acetylation of α-tubulin and the axonal transport defects in flies and mice that lack Elongator activity

In order to test whether Acly is required for proper axonal transport in fly, we generated *Acly* KD flies (42) that displayed reduced α-tubulin acetylation (Figure 3a) together with impairment of axonal transport of synaptotagmin (SYT1-GFP) vesicles (Figures 3b-d, Figure S3a). In agreement with our observation made with Elp3 cKO mice (Figures 2m, 2n), the expression of Acly protein was also reduced in the brain of adult Elp3 RNAi flies (Elav:GAL4; Elp3RNAi) (Figure 3e). We postulated that the acetylation of α-tubulin and the axonal transport defects observed upon loss of Elongator may arise from a reduced expression of Acly. To test this hypothesis, we expressed human ACLY (which shares high amino acid homology with its murine and fly homologues; Figure S3b) in *Elp3* KD larva MNs (UAS:Elp3 RNAi and UAS:ACLY, *Elp3* KD+ACLY) and in *Elp3* cKO mouse cortical PNs and showed that it raised the acetylation of α-tubulin in both flies MNs *in vivo* (Figure 3f) and in culture mouse cortical PNs *in vitro* (Figure 3l). Those data were correlated with a rescue of all axonal transport parameters, including the average and moving bi-directional transport velocities together with pausing time of SYT1-GFP vesicle in MNs of 3^rd^ instar larvae (Figures 3g-i, Figure S3c) as well as of lysosomes and mitochondria in cultured PNs from *Elp3* cKO mice (Figures 3m-o, Figure S3d-g). Furthermore, the locomotion activities of both *Elp3* KD 3^rd^ instar larvae and adult flies were restored to control levels upon ACLY expression (Figure 3j-k). Altogether, these results suggest that the reduction of Acly expression leads to MTs acetylation and axonal transport defects in Elongator deficient neurons.

**Figure 3.**
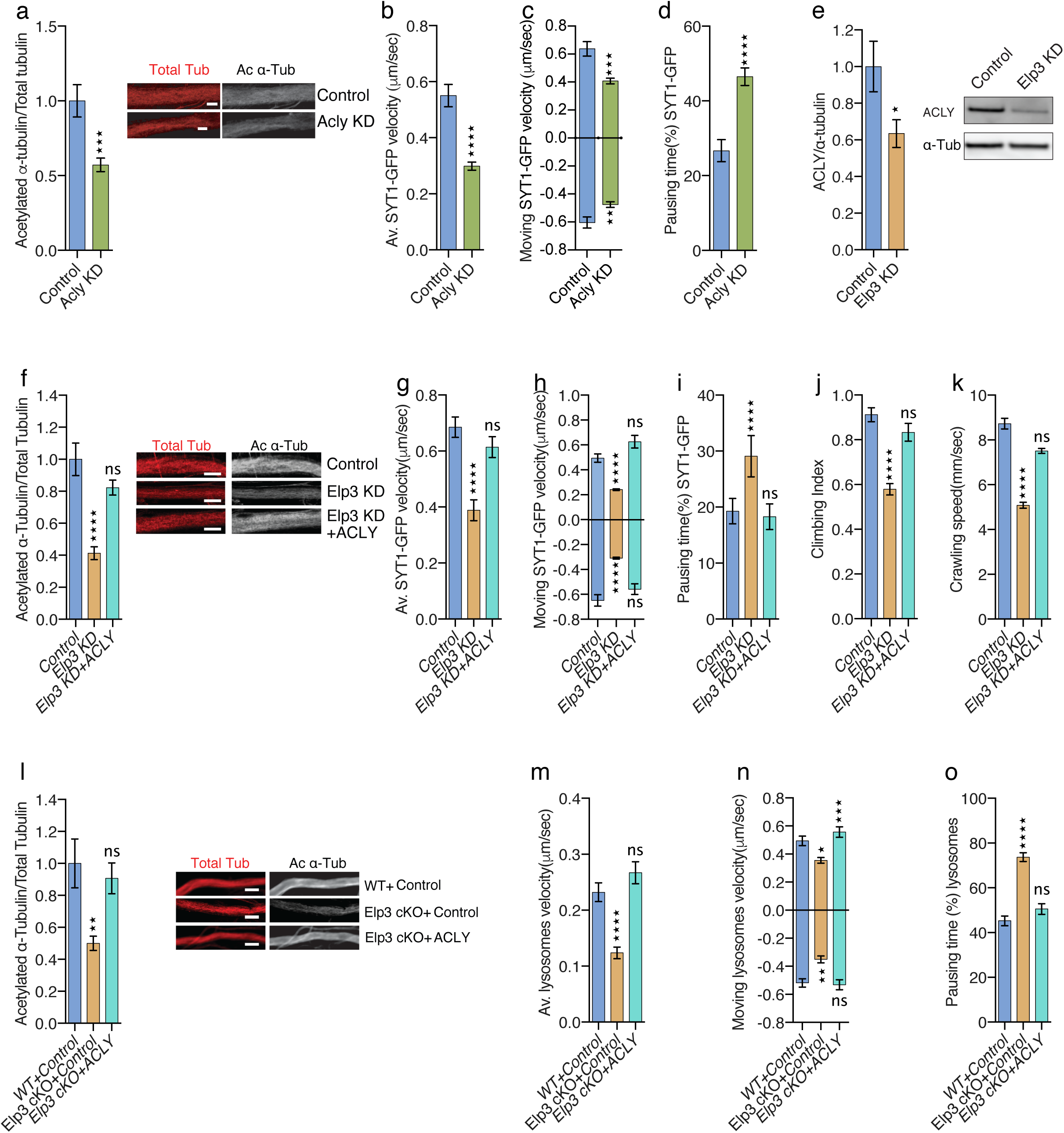
ACLY expression rescues α-tubulin acetylation and molecular transport defects upon loss of Elongator activity in mouse and fly neurons. (**a**) Immunolabelings and quantification of acetylated α-tubulin (Ac α-Tub) and total tubulin (Tot Tub) in 3^rd^ instar larvae of *Drosophila melanogaster* control or expressing Acly RNAi (Acly KD) under a motoneuron-specific driver (D42:GAL4). Scale bar is 10 μm. (**b-d**) Time-lapse recording of Synaptotagmin-GFP (Syt1-GFP) axonal transport in 3^rd^ instar larvae motor neurons of control or *Acly* KD to analyze average (av.) velocity (**b**), moving velocity (**c**) and percentage of pausing time (**d**). (**e**) Immunoblotting and quantification of Acly and total tubulin expressions in *Drosophila melanogaster* head extracts from control or Elp3 RNAi (Elp3 KD) (under the pan-neuronal driver, Elav:GAL4) adult flies. (**f**) Immunolabelings and quantification of acetylated α-tubulin (Ac α-Tub) and total tubulin (Tot Tub) in motoneuron axons of 3^rd^ instar larvae: control, *Elp3* KD and *Elp3* KD + human *ACLY*. Scale bar is 10 μm. (**g-k**) *In vivo* live imaging and behavior measurements in 3^rd^ instar larva: control, *Elp3* KD and *Elp3* KD + human *ACLY*. (**g-i**) Time-lapse recording of axonal transport of Synaptotagmin-GFP (Syt1-GFP) in motoneurons to analyze average (av.) velocity (**g**), moving velocity (**h**), and percentage of pausing time (**i**). (**j-k**) Locomotion assays; histograms of climbing index of adult flies (**j**) and crawling speed of 3^rd^ instar larvae (**k**). (**l**) Immunolabelings (axons are shown) and quantification of acetylated α-tubulin (Ac α-Tub) and total tubulin (Tot Tub) in cultured cortical PNs from E14.5 WT and *Elp3cKO* embryos transfected with ACLY or control. Scale bar is 10 μm. (**m-o**) cortical neurons isolated from E14.5 WT and Elp3cKO embryos were transfected with control or ACLY expressing constructs and cultured for 5 DIV in microfluidic devices to perform time-lapse recording of axonal transport and measure average (av.) velocity (**m**), moving velocity (**n**) and percentage of pausing time (**o**) of lysosomes (LysoTracker®). Description of graphical summaries here within are histograms of means ± SEM, while statistical analyses of (**a**,**e**) are two-tailed t-test, (b, **c, d**) are Mann-Whitney test, (**f, h, j, k, l**) are one-way analysis of variance (ANOVA), (**g, i, m, n, o**) is Kruskal Wallis one-way ANOVA. Specifically, [(**a**) p=0.0004, t=3.690, df=99; (**b**) p<0.0001, U= 24389; (**c**) p=0.0007, U=3453 and p=0.0072, U=4945 for anterograde and retrograde, respectively; (**d**) p<0.0001, U= 30545; (**e**) p=0.031, t=2.407, df=13; (**f**) p=0.0003, F=9.158; (**g**) p<0.0001, K=21.66; (**h**) p<0.0001, F=25.90 and p<0.0001, F=20.96 for anterograde and retrograde, respectively; (**i**) p<0.0001, K=22.35; (**j**) p<0.0001, F=28.45; (**k**) p<0.0001, F=139.5; (**l**) p=0.0042, F=4.969; (**m**) p<0.0001, K=28.69; (**n**) p=0.001, K=13.85 and p<0.0001, K=251.5; (**o**) p<0.0001, K=24.74. In addition, post hoc multiple comparisons for (**f**) is Dunnet’s test, for (**h**) is Holm-Sidak’s test, for (**g**,**i**) are Dunn’s test, for (**j**,**k**,**l**,**m**,**n**,**o**) are Sidak’s test and are *p<0.05, **p < 0.01, ***p < 0.001, and ****p < 0.0001. The total number of samples analyzed were as follows: (**a**) 50 to 57 motoneurons from at least 6 larvae per group; (**b-d**) 170-421 vesicles coming from 12 different larvae per group (at least 3 images per animal); (**e**) 7-8 flies per group; (**f**) 12 to 37 neurons from at least 3 different mice per group; (**g-i**) 78-304 vesicles tracks from 5-10 larvae per group; (**j**) 8-9 test per group containing 10 adult flies; (**k**) 11 to 15 larvae per group, (**l**) 23-28 neurons coming from 3-5 different mice per group; (**m-o**) 343 to 443 vesicles tracks coming from 3-5 different mice per group (at least 5 images per mouse).

### Fibroblasts from Familial Dysautonomia patients show defects of tubulin acetylation and MT-dependent transport

In order to place our findings in a human pathological context, we analyzed primary skin fibroblasts from FD patients. These cells were isolated from FD patients carrying the splice site IVS20+6T_C variant in *ELP1* (43) that expressed barely detectable amount of ELP1 proteins and low level of ELP3 that results from its instability upon non-assembly of the Elongator complex (Figure S4a) (25). We observed a reduction of acetylation of α-tubulin and MT-dependent transport defects in FD fibroblasts (Figure S4b-f), which were comparable to those observed in fly and mouse neurons. This modification was not resulting from an increased deacetylation activity of HDAC6 towards MTs in extracts fibroblasts nor a massive intra-cellular protein aggregation that would affect MT-dependent transport in FD fibroblasts (Figures S4g-s4h). Both α-tubulin acetylation and MT-dependent transport parameters were corrected in FD fibroblasts by blocking tubulin deacetylation with incubation of the HDAC6 specific inhibitor, Tubastatin (TBA; 10uM during 30 min) (Figure S4b). Similarly to the animal models, FD fibroblasts express reduced amount of ACLY proteins (Figures 4a) and incubation with the ACLY competitive inhibitor Hydroxycitrate (HCA) for 8 hours interfered in a dose-dependent manner with MT acetylation (Figure 4b), in both WT and FD fibroblasts and in line with the residual expression of ACLY observed in FD fibroblasts (Figure 4a). We next showed that overexpression of ACLY in FD fibroblasts rescued the level of acetylation of α-tubulin (Figure 4c) as well as the defects of MT-dependent transport of lysosomes (Figures 4d-4f, S4i). Altogether, these results demonstrate that a reduction of ACLY in FD fibroblasts underlies MT acetylation and MT-dependent transport impairments; these molecular defects likely contribute to the pathological mechanisms underlying FD.

**Figure 4.**
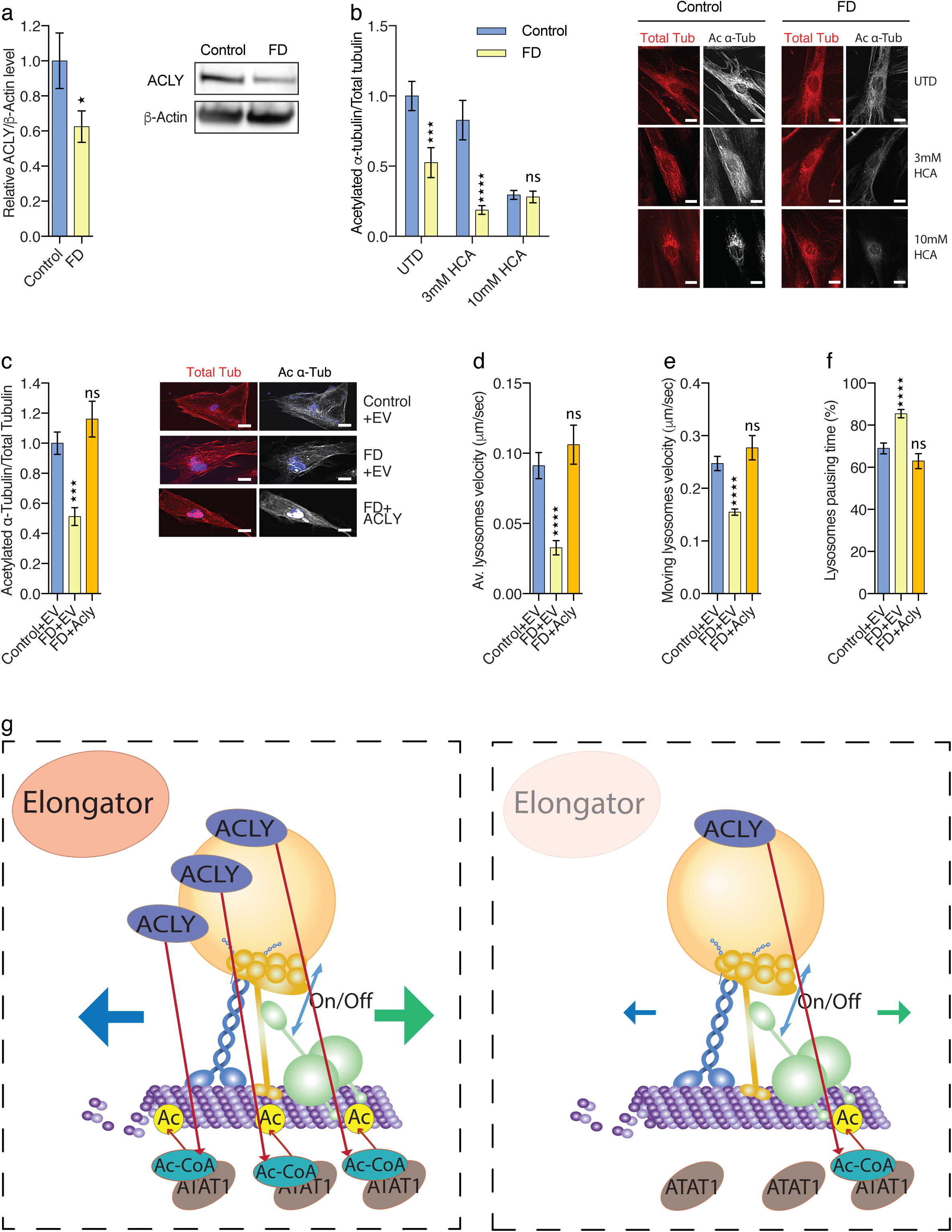
ACLY expression rescues defects of α-tubulin acetylation and microtubule-dependent transport in human FD fibroblasts. (**a**) Immunoblotting of ACLY and ß-ACTIN in human primary fibroblast extracts from Control and FD patients. (**b**) Immunolabelings and quantification of acetylated α-tubulin (Ac α-Tub) and total tubulin (Tot Tub) in primary fibroblasts from healthy controls and FD patients incubated with ACLY inhibitor hydroxycitrate (HCA). Scale bar is 50 μm. (**c**) Immunolabelings and quantification of acetylated α-tubulin and (Ac α-Tub) total tubulin (Tot Tub) in extracts from primary fibroblasts of healthy controls and FD patients transfected with control or ACLY expressing plasmids. Scale bar is 50 μm. (**d-f**) Time-lapse recording of intra-cellular lysosome transport (LysoTracker®) in fibroblasts from Control or FD patients, transfected with Control or ACLY plasmids, to analyze average (av.) velocity (**d**), moving velocity (**e**) and percentage of pausing time (**f**). (**g**) Summary scheme showing a central role played by Acly in both the control of α-tubulin acetylation and microtubule-dependent transport, which is impaired upon loss of Elongator activity. Description of graphical summaries here within are histograms of means ± SEM, while statistical analyses of (**a**) is two-tailed t-test, (**b**) is two-way ANOVA, (**c**) is one-way analysis of variance (ANOVA), (**d, e, f**) are Kruskal Wallis one-way ANOVA. Specifically, [(**a**) p= 0.0467, t=2.181,df=14; (**b**) p=0.0038, F_interaction_ (2, 109) = 5.876; (**c**) p<0.0001, F=13.17; (**d**) p<0.0001, K= 48.95; (**e**) p<0.0001, K=35.10; (**f**) p<0.0001, K=40.98. In addition, post hoc multiple comparisons for (**b**) is Sidak’s test, for (**c**) is Dunnet’s test, for (**d, e, f**) are Dunn’s test, and are *p<0.05, ***p < 0.001, and ****p < 0.0001. The total number of samples analyzed were as follows: (**a**) 5 human primary fibroblasts per group; (**b, c**) 15 to 26 human primary fibroblasts from 4-5 human primary fibroblasts per group; (**d-f**) 15 images per group from 5 human primary fibroblasts per group; 143-217 vesicles tracks from human fibroblast cells from 5 human primary fibroblasts per group (at least 5 images per sample).

## Discussion

Here we show that loss of Elongator activity impairs both MT acetylation and bidirectional axonal transport in mouse cortical neurons in culture as well as in fly motoneurons *in vivo*, ultimately resulting in locomotion deficits in flies. These defects are similar to those observed upon loss of Atat1 expression, the enzyme that catalyzes the acetylation of α-tubulin K40 residues in MTs. This shared phenotype prompted us to investigate a possible collaboration between Elongator and Atat1 in the control of MT-dependent axonal transport via *α*-tubulin acetylation. By combining conditional loss of function models with genetic complementation experiments, we observed a molecular cooperation between these molecules and identified Acly as the common denominator, which provides acetyl groups from Acetyl-CoA to vesicular Atat1 and whose expression depends on Elongator activity. We showed that raising Acly level not only rescued axonal transport defects in both mouse and fly Elongator models but also improved locomotion activity of *Elp3* KD larvae and flies. Moreover, loss of Elongator activity does not affect *Acly* transcription but prevents the accumulation of its proteins in neurons. Since the expression of ACLY rescues defects of α-tubulin acetylation and axonal transport in both mouse and fly Elongator models, it is unlikely the reduced amount of Acly proteins detected by western blotting mainly results from the poor translation of its corresponding transcripts. However, Elongator may promote Acly protein stabilization via a direct interaction, as previously reported for other cytoplasmic proteins. Another non-mutually exclusive mechanism that may act downstream Elongator would involve the regulation of the acetyltransferase (44) P300/CBP-associated factor (PCAF), which is known to increase Acly stability by promoting its acetylation (44). Indeed, the molecular mechanism linking Elongator to Acly expression deserves more attention and is currently under investigation in the laboratory.

Elongator subunits and Atat1 have been detected in protein extracts from purified motile vesicles (16, 30) and the present study confirms those data and shows that a functional pool of Acly is also predominantly enriched in the vesicular fraction of mouse cerebral cortical extracts. Therefore, we postulate that an Elongator/Acly/Atat1 (EAA) signaling pathway may directly act at vesicles to promote MT acetylation, further supporting their non-canonical role as a platform for local signaling and for regulating axonal transport in particular (Figure 4g). Since these molecules are also detected in the cytosol (but only a small pool for Atat1 and Acly; Figure 2k), we cannot rule out a minor contribution of a cytosolic EAA pathway to these processes. We observed comparable defects of *α*-tubulin acetylation and MT-dependent transport in fly, mouse, and human cells lacking Elongator activity, which, together with similar observations made in C. elegans (6), suggest a strong functional conservation of the EAA pathway across species.

A tight control of axonal transport is indeed very important to ensure homeostatic activity to neurons by delivering cargos to distant regions, thereby controlling cytoskeleton maintenance as well as spreading long range intracellular signaling that ensures cellular maintenance and function. Impairment of axonal transport is indeed considered as an early pathological feature shared by several neurodegenerative disorders (2, 45, 46). Along this line, our results obtained with FD fibroblasts support those previously described in neurons (28) and suggest that loss of Elongator activity contribute to neurodegeneration by interfering with MT-dependent transport in FD patients via impairment of ACLY expression. We did not observe differences in MT deacetylation activity in FD and WT fibroblasts (Figures S4g), suggesting that reduction of MT-acetylation in FD fibroblasts does not arise from changes of HDAC6 expression, in contrast to what others have reported (28).

More generally, *α*-tubulin acetylation not only modulates axonal transport but also provides resistance to mechanical bending to MTs (47). Therefore, by controlling the cytoskeleton integrity and function, the EAA pathway likely acts as a key regulator of neuronal fitness whose progressive dysfunction may contribute to neuronal aging and even underlies neurodegeneration. Thus, targeting this molecular pathway may open new therapeutic perspectives to prevent the onset or the progression of neurodegenerative disorders characterized by poor axonal transport and degeneration.

**Figure S1.**
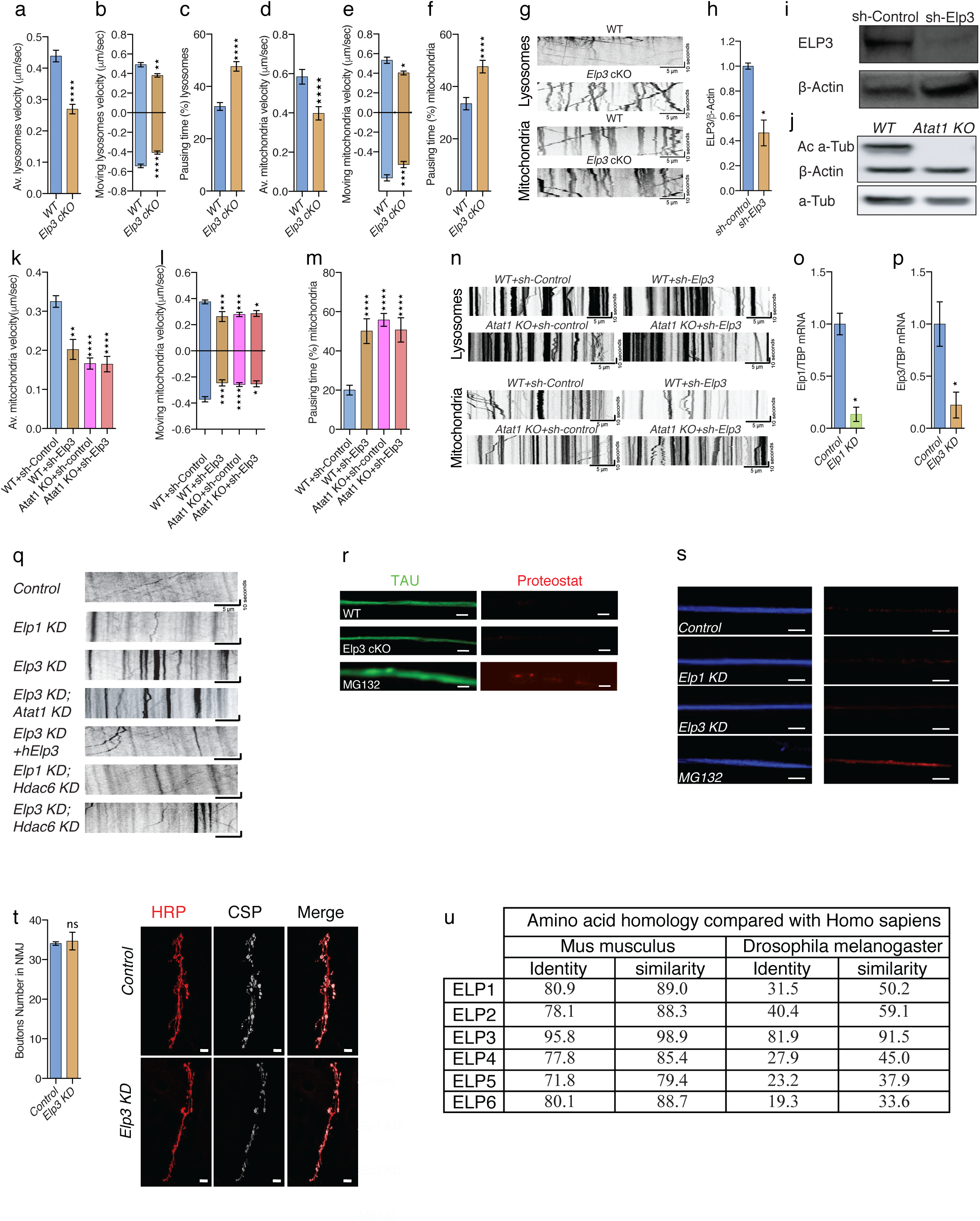
Kymographs of motile lysosomes, mitochondria and Synaptotagmin-GFP. RNAi targeting efficiency and neuromuscular junction assessment in fly model. (**a-f**) Axonal transport recordings in E14.5 mice cortical neurons cultured for 5 DIV in microfluidic devices for time-lapse recording of axonal transport from WT or *Elp3 cKO* neurons to analyze average (av.) velocity (**a, d**), moving velocity (**b, e**) and percentage of pausing time (**c, f**) of lysosomes (LysoTracker®) and mitochondria (MitoTracker®). (**g**) Representative kymographs of Lysosomes (**top**) and Mitochondria (**bottom**) in E14.5 in *WT or Elp3cKO* mice cortical neurons axons cultured for 5 DIV in microfluidic devices. Scale bars are 10 seconds and 5 μm. (**h-i**) Immunoblotting of Elp3 and ß-Actin in cortical neurons transfected with sh-*Elp3* or sh-*control*. (**j**) Immunoblotting of Acetylated α tubulin (Ac α-Tub), Total α tubulin (Tot α-Tub) and ß-Actin in E14.5 WT and *Atat1 KO* mice brain cortex. (**k-m**) Axonal transport study of mitochondria (MitoTracker®) in WT or *ATAT1 KO* neurons infected either with sh-Control or sh-*Elp3* to analyze average (av.) velocity (**k**), moving velocity (**l**) and percentage of pausing time (**m**). (**n**) Representative kymographs of Lysosomes (**top**) and Mitochondria (**bottom**) in E14 in WT or *ATAT1 KO* neurons infected either with sh-Control or sh-*Elp3*mice cortical neurons axons cultured for 5 DIV in microfluidic devices. Scale bars are 10 seconds and 5 μm. (**o-p**) qRT-PCR analysis to measure mRNA levels of *Elp1* and *Elp3* from fly head extracts, expressing RNAi under the pan-neuronal driver (Elav:GAL4) in control, *Elp1* KD (**o**) and *Elp3* KD (**p**). (**q**) Representative kymographs of Synaptotagmin-GFP (Syt1-GFP) in motoneurons from 3^rd^ instar larvae of *Drosophila melanogaster* expressing RNAi under a motoneuron-specific driver (D42:GAL4); control, *Elp1* KD, *Elp3* KD, *ATAT1; Elp3* KD, *Elp3* KD + human *Elp3, Elp1;Hdac6* KD and *Elp3;Hdac6* KD. Scale bars are 10 seconds and 5 μm. (**r**) Protein aggregation was estimated in axons (Tau+; in green) of cultured mice cortical neurons of WT, *Elp3cKO* and MG132 treated neurons (positive control) by the chemical dye Proteostat© as marker for protein aggregation (in red). Scale bar is 10 μm. (**s**) Protein aggregation was estimated in motor neurons exiting the ventral ganglion of 3^rd^ instar larvae expressing RNAi under motoneurons specific driver (D42:GAL4); control, *Elp1* KD, *Elp3* KD and *MG132 treatment* (positive control) by immunolabeling of cysteine string protein(CSP+; red) as marker for aggregation and the neuronal marker horseradish peroxidase (HRP+; blue). Scale bar is 10 μm. (**t**) Comparison of Elongator complex subunits (ELP1-6) amino acid homology from mice and flies with human by sequence identity and similarity. (**u**) Representative images (**right**) of neuromuscular junctions in 3^rd^ instar larvae immunolabeled for cysteine string protein (CSP) and horseradish peroxidase (HRP) and a histogram of its quantification (**left**). Description of graphical summaries here within are histograms of means ± SEM, while statistical analyses of (**a, b, c, d, e, f, h, o, p, u**) are Mann-Whitney test, (**k, l, m**) are Kruskal Wallis one-way ANOVA. Specifically, [(**a**) p<0.0001, U=111906; (**b**) p<0.0001, U=71910 and p=0.0029, U=45529 for anterograde and retrograde, respectively ; (**c**) p<0.0001, U=68287; (**d**) p<0.0001, U=21672; (**e**) p=0.0349, U=20092 and p=0.0002, U=27975 for anterograde and retrograde, respectively; (**f**) p<0.0001, U= 21907; (**h**) p=0.0286, U=0 ; (**k**) p<0.0001, K=67.88; (**l**) p<0.0001, K=25.42 and p<0.0001, K=25.89 for anterograde and retrograde, respectively; (**m**) p<0.0001, K=73.34; (**o**) p=0.0159, U=0; (**p**) p=0.0159, U=0; (**u**) p=0.6905, U=10]. In addition, post hoc multiple comparisons for (**k, l, m**) are Dunn’s test, and are *p<0.05,**p<0.01, ***p < 0.001, and ****p < 0.0001. The total number of samples analyzed were as follows: (**a, c, e**) 423 to 712 vesicles tracks from 5 different mice per group; (**b, d, f**) 232 to 239 vesicles tracks from 5 different mice per group; (**h**) 4 animals per group; (**k, l, m**) 83 to 201 vesicles tracks from 3 different mice per group; (**o, p**) 4 to 5 fly’s brain per group; (**u**) 5 3^rd^ instar larvae per group (average of 3-5 NMJs each).

**Figure S2.**
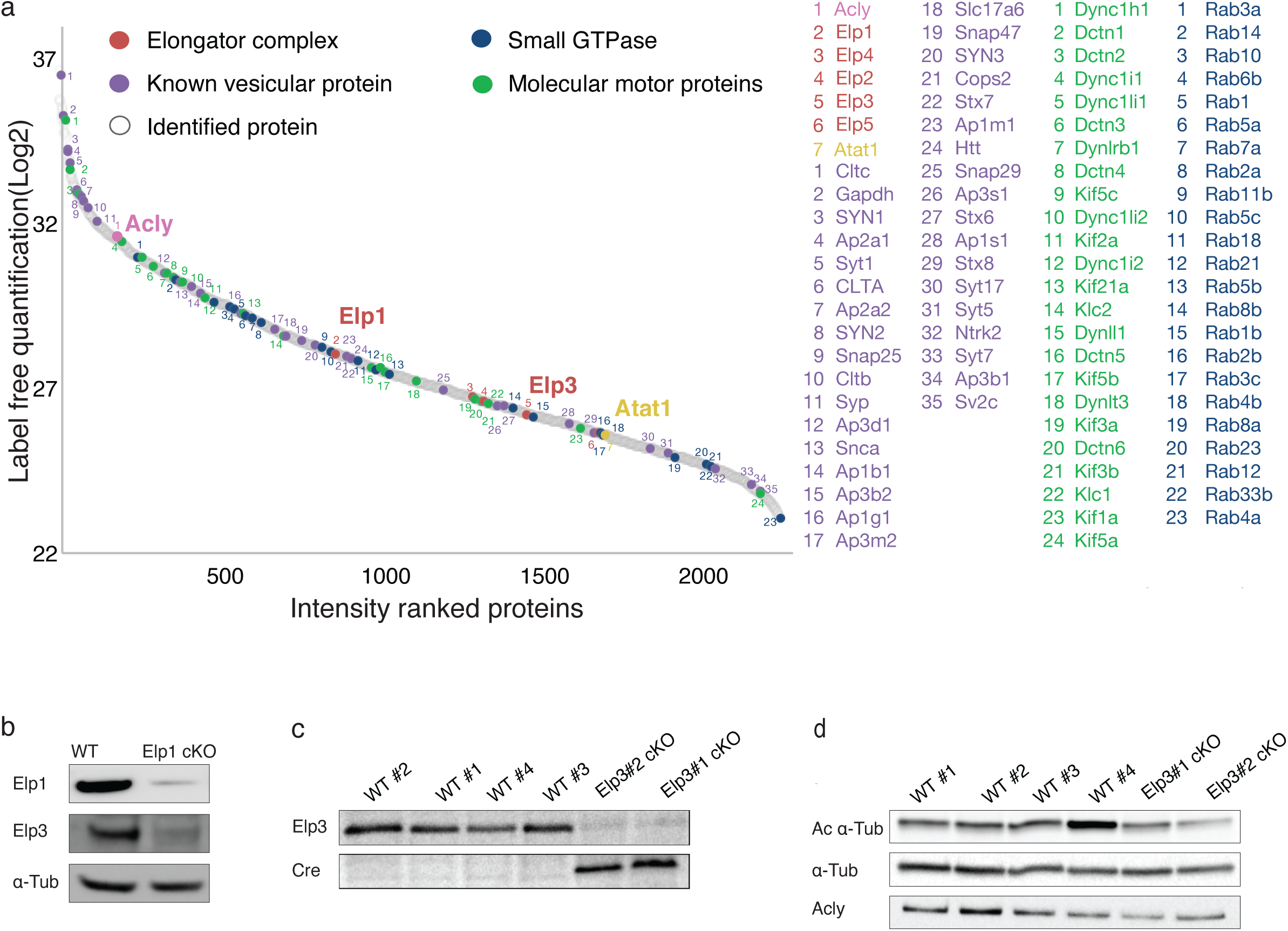
LC-MS/MS and Acly levels in E14.5 cortical brains. (**a**) LC-MS/MS proteomic analysis of vesicular fraction isolated from adult brain cortices of WT and Elp1 cKO mice, proteins were ranked by intensity and plotted according to their relative abundance (gray spots). Acly (pink), Atat1 (yellow), and Elongator subunits (red) detection among proteins previously identified as vesicular components (purple), small GTpase (blue) and molecular motors (green) (n = 3, graph represent the mean intensity value). (**b**) Immunoblotting to detect Elp1, Elp3 and α-tubulin (α-Tub) in cortical extracts from adult WT and *Elp1 cKO* mice. (**c**) Immunoblotting to detect Elp3 and Cre. (**d**) Immunoblotting to detect Elp3, Ac α-Tubulin, α-Tubulin (α-Tub) and Acly in cortical extracts from E14.5 WT and *Elp3 cKO* mice (n=2 to 3 E14.5 mouse embryos).

**Figure S3.**
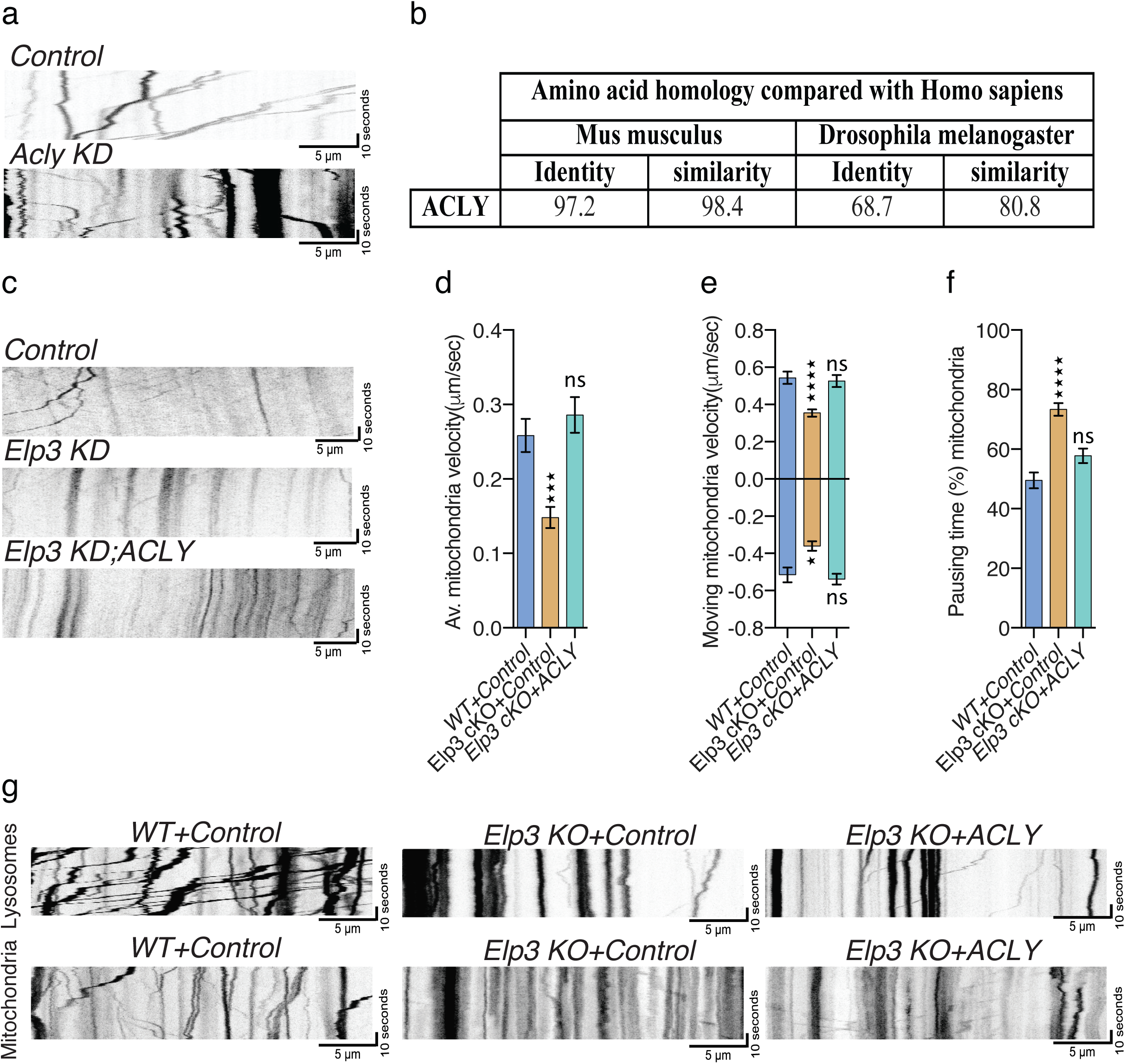
Expression of ACLY in mice cultured neurons and fly heads. Kymographs after time-lapse recording of motile lysosomes, mitochondria and Synaptotagmin-GFP+ vesicles. (**a**) Representative kymographs of Synaptotagmin-GFP (Syt1-GFP) in motoneurons from 3^rd^ instar larvae of *Drosophila melanogaster* expressing RNAi under a motoneuron-specific driver (D42:GAL4); control and *Acly* KD. Scale bars are 10 seconds and 5 μm. (**b**) Comparison of Acly amino acid homology from mice and flies with human by sequence identity and similarity. Representative kymographs of Synaptotagmin-GFP (Syt1-GFP) in 3^rd^ instar larvae motor neurons expressing RNAi under a motoneuron-specific driver (D42:GAL4); control, *Elp3* KD and *Elp3* KD + *ACLY*. Scale bars are 10 seconds and 5 μm. (**d-f**) E14.5 mice cortical neurons cultured for 5 DIV in microfluidic devices for time-lapse recording of axonal transport of mitochondria (MitoTracker®) from *WT or Elp3cKO* neurons transfected with control or *Acly* expressing constructs, to analyze average (av.) velocity (**d**), moving velocity (**e**) and percentage of pausing time (**f**). (**g**) Representative kymographs of Lysosomes (**top**) and Mitochondria (**bottom**) in E14.5 in *WT or Elp3cKO* mice cortical neurons axons transfected with ACLY or Control carrying construct and cultured for 5 DIV in microfluidic devices. Scale bars are 10 seconds and 5 μm. Description of graphical summaries here within are histograms of means ± SEM, while statistical analyses of (**d, e, f**) are Kruskal Wallis one-way ANOVA. Specifically, [(**d**) p<0.0001, K=27.47; (**e**) p<0.0001, K=26.16 and p=0.0003, K=16.04 for anterograde and retrograde, respectively; (**f**) p<0.0001, K=49.32]. In addition, post hoc multiple comparisons for (**d, e, f**) are Dunn’s test, and are *p<0.05, ***p < 0.001, and ****p < 0.0001. The total number of samples analyzed were as follows: (**d, e, f**) 283 to 340 vesicles tracks from 5 different mice per group.

**Figure S4.**
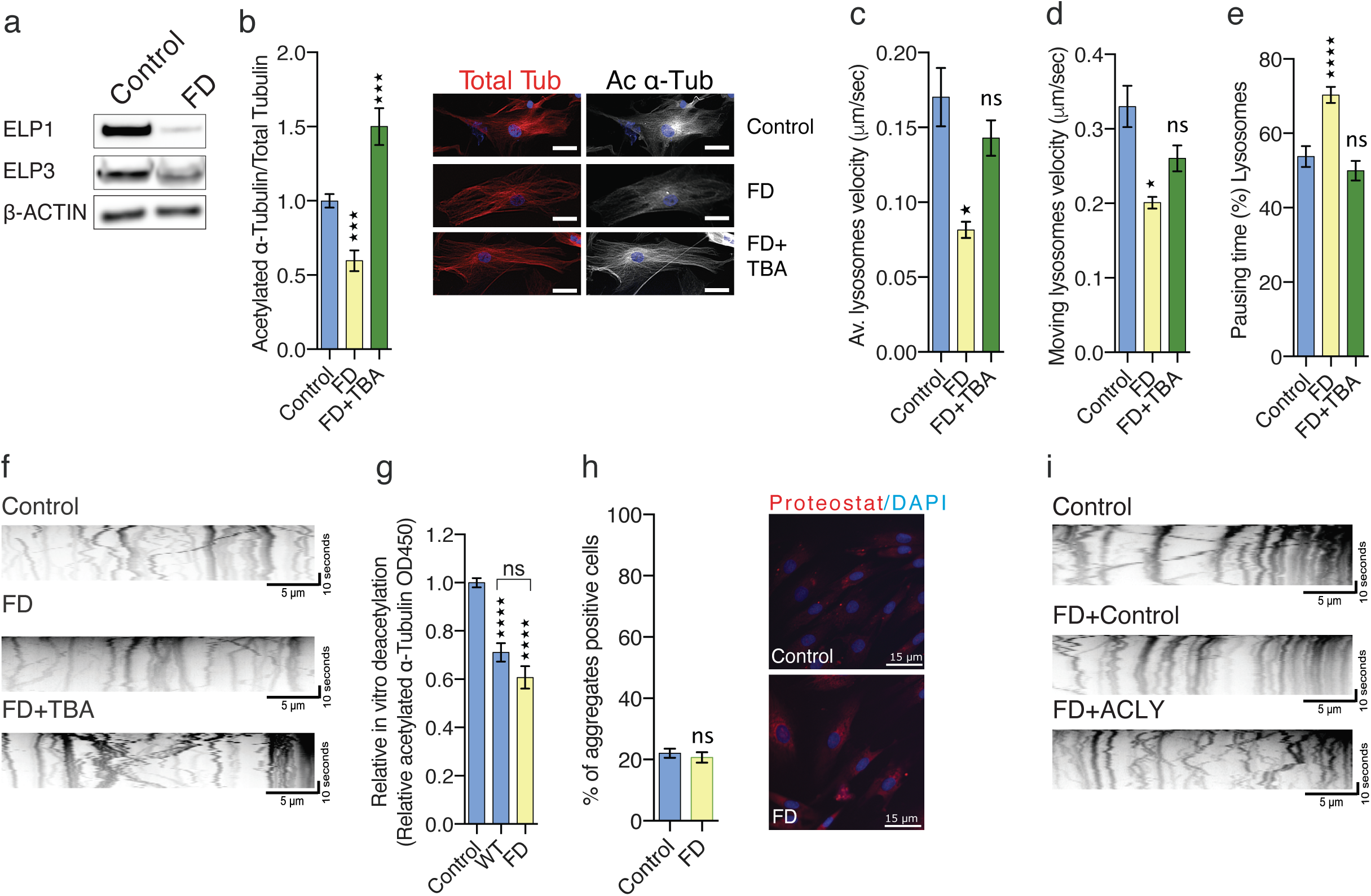
Pharmacological and genetic rescue of vesicular transport in FD fibroblasts. Kymographs form time-lapse recorded motile lysosomes in human primary fibroblasts. **(a)** Immunoblotting of ELP1, ELP3 and ß-ACTIN in human primary fibroblasts from control and FD patients. (**b**) Immunolabelings and quantification of acetylated α-tubulin (Ac α-Tub) and total tubulin (Tot Tub) in human primary fibroblasts from healthy controls and FD patients incubated with vehicle or Tubastatin (TBA). Scale bar is 50 μm. (**c-e**) Detection of lysosomes using live fluorescent probe (LysoTracker®) in human primary fibroblasts for time-lapse recording of intracellular transport in control or FD cultured fibroblasts incubated with vehicle (DMEM media) or TBA to analyze average (av.) velocity (**c**), moving velocity (**d**) and percentage of pausing time (**e**). (**f**) Representative kymographs of transported lysosomes in human primary fibroblasts from healthy controls or FD patients incubated with vehicle (DMEM media) or Tubastatin (TBA). Scale bars are 10 seconds and 5μm. (**g**) In vitro deacetylation assay of endogenously acetylated bovine brain tubulin incubated for 4 hours with extracts of control and FD cultured fibroblasts or without tissue extract (control). (**h**) Protein aggregation was estimated in fibroblasts from healthy controls and FD patients after incubation with the chemical dye Proteostat©, a marker for protein aggregation (in red). Scale bar is 50 μm. (**i**) Representative kymographs of transported lysosomes in human primary fibroblasts from healthy controls or FD patients transfected with ACLY or Control carrying constructs. Scale bars are 10 seconds and 5μm. Description of graphical summaries here within are histograms of means ± SEM, while statistical analyses of (**b**) one-way analysis of variance (ANOVA), (**c, d, e**) are Kruskal Wallis one-way ANOVA (**g**) is one-way ANOVA and (**h**) is Mann-Whitney test. Specifically, [(**b**) p<0.0001, F=30.36; (**c**) p<0.0001, K=24.26; (**d**) p<0.0001, K=23.82; (**e**) p<0.0001, K=44.79; (**g**) p=0.3398, U=60]. In addition, post hoc multiple comparisons for (**b**) is Dunnett’s test, for (**c, d, e**) are Dunn’s test, and are *p<0.05, ***p < 0.001, and ****p < 0.0001. The total number of samples analyzed were as follows: (**a**) lysates from five independent cultures of control and FD patient fibroblasts; (**b**) average of 13 to 30 fibroblasts neurons from 5 human primary fibroblasts from control and FD patients (3 technical replicates each). (**c, d, e**) 225 to 273 vesicles tracks from 5 different human primary fibroblasts from control and FD patients (3 technical replicates each); (**g**) 4-5 extracts 5 of human primary fibroblasts from control and FD patients (**h**) average of 12 to 13 fibroblasts from 5 human primary fibroblasts of control and FD patients (3 technical replicates each).

## Material and Methods

### Mice

Brains were harvested from mice at P0-P2 or E14.5. *FoxG1:*Cre ^-/+^/*Elp3* ^lox/+^ and *Elp3* ^*l*ox/lox^ mice were time-mated to induce a loss of function of *Elp3* in the forebrain (48). Brains were harvested from 10-16 months-old Elp1cKO and WT mice (27).

*Atat1* ^+/-^ mice were bred to obtain WT and KO mice (49). Mice were housed under standard conditions and they were treated according to the guidelines of the Belgian Ministry of Agriculture in agreement with the European Community Laboratory Animal Care and Use Regulations (86/609/CEE, Journal Official des Communautés Européennes L358, 18 December 1986).

Neuronal cultures were prepared from dissected E14.5 mice brain cortices, followed by mechanical dissociation in HBBS (Sigma-Aldrich, H6648) supplemented with 1.5% glucose. Cells were cultured at confluence of ∼70% with Neurobasal Medium (Gibco, Invitrogen, 21103049) supplemented with 2% B27 (Gibco, Invitrogen, 17504044), 1% Pen/Strep (Gibco, Invitrogen, 15140122) and 1% Glutamax (Gibco, Invitrogen, 35050061) at 37°C.

### Drosophila melanogaster

Flies were kept in 25°C incubator with regular 12-hour light and dark cycle. All crosses were performed at 25°C, after two days hatched first instar larvae were transferred in a 29°C incubator until use.

UAS-RNAi carrying lines were crossed with virgin females were crossed with of D42-Gal4-UAS:Syt1-GFP for axonal transport recording, or with D42-Gal4 for behavioral experiments and immuno-staining, or with Elav-Gal4 for qPCR and WB analysis. All RNAi inserts sequences were validated by DNA sequencing.

*Drosophila melanogaster* lines:

| Line | Catalog number | Given Name | Purchased from |
| --- | --- | --- | --- |
| Elav-Gal4 | BDSC 458 |  | Bloomington Drosophila Stock Center |
| huD42-Gal4 | BDSC 8816 |  | Bloomington Drosophila Stock Center |
| UAS:Syt-GFP | BDSC 6925 |  | Bloomington Drosophila Stock Center |
| UAS:RNAi Zpg | VDRC CG10125 | Control | Vienna Drosophila Resource Center |
| UAS:RNAi Elp1 | VDRC CG10535 | Elp1 KD | Vienna Drosophila Resource Center |
| UAS:RNAi Elp3 | VDRC CG15433 | Elp3 KD | Vienna Drosophila Resource Center |
| UAS:RNAi Atat1 | VDRC CG3967 | Atat1 KD | Vienna Drosophila Resource Center |
| UAS:RNAi Hdac6 | BDSC 51181 | Hdac6 KD | Bloomington Drosophila Stock Center |
| UAS:Human Elp3 |  | hElp3 | Kindly provided by Patrick Verstrecken |
| UAS:Human ACLY | VDRC CG8322 | hACLY | Vienna Drosophila Resource Center |

## Human primary fibroblasts

Fibroblasts were cultured in polystyrene culture flasks (Corning) at 37°C with 5% CO2 in DMEM (Gibco, Invitrogen, 11965092) medium supplemented with 10% Fetal Calf Serum (Gibco, Invitrogen, 10500056), 1 mM Sodium pyruvate (Gibco, Invitrogen, 11360070), 1 mM non-essential amino acids (Gibco, Invitrogen, 11140050). Primary fibroblasts from five FD patients and age matched controls were purchased from Coriell biobank (www.coriell.org).

| Catalog number | Sex | Age | Sample Description |
| --- | --- | --- | --- |
| GM02036 | Female | 11 | Apparently Healthy |
| GM07492 | Male | 17 | Apparently Healthy |
| GM07522 | Female | 19 | Apparently Healthy |
| GM038 | Female | 3 | Apparently Healthy |
| GM05659 | Male | 14 | Apparently Healthy |
| GM02343 | Female | 24 | Familial Dysautonomia |
| GM04589 | Male | 16 | Familial Dysautonomia |
| GM04663 | Female | 2 | Familial Dysautonomia |
| GM04959 | Female | 10 | Familial Dysautonomia |
| GM04899 | Female | 12 | Familial Dysautonomia |

### Immunofluorescence

E14.5 cortical neurons were plated on coverslips in 24 well plate at density of 30,000 cells and cultured for 5DIV. Neurons were fixed by incubation in 4% PFA in PBS for 20 minutes at RT and washed with PBS+0.3% Triton-X. After washing, neurons were incubated in blocking solution (PBS+0.3% Triton-X+10% normal donkey serum) for 1 hour at RT. Following overnight incubation with primary antibodies in blocking solution at 4°C, washing, and incubation with secondary antibodies (PBS+0.3% Triton-X+1% normal donkey serum) at RT for 1 hour and washing, coverslips were then mounted on microscope slide using Mowiol. Images were acquired using Nikon, A1Ti, confocal microscope, 60X lens or Airyscan super-resolution module on a Zeiss LSM-880 confocal.

Larvae were dissected in PBS to expose brain and motor neurons, after dissection larvae were fixed with 4% PFA for 20 minutes at RT, washed with PBS+0.2% (CSP staining) or 0.3% (*α*-Tubulin acetylation staining) Triton-X and incubated in blocking solution PBS, 0.2% (CSP staining) or 0.3% (*α*-Tubulin acetylation staining) Triton-X+1% BSA for 30 minutes at RT. Following overnight incubation with primary antibodies at 4°C, washing and incubation with secondary antibodies at RT for 2 hours the larvae were mounted on microscope slide in Mowiol. Images were acquired using Nikon, A1Ti, confocal microscope, 60X lens or Airyscan super-resolution module on a Zeiss LSM-880 confocal.

Primary fibroblasts from FD patients or controls were plated at concentration of 5,000 cells in 96 well plate (uClear, Grinere). Cells were fixed with 4% PFA for 10 minutes, washed three times with PBS, permeabilized and blocked for 1 hour at RT with 5% fetal bovine serum in PBS+0.1% Triton-X. Following overnight incubation with acetylated α-tubulin (Sigma-Aldrich) and <u>β</u>-tubulin (cell signaling) antibodies, cells were washed three times with PBS+0.05% Triton-X times and incubated with secondary antibodies for 1hour at RT. Finally, the cells were washed three times, and remained in PBS for image acquisition. Images were acquired using in-cell 2200 (General Electric) fluorescent microscope using 60X air lens.

Fluorescence intensity levels were measured by Fiji (https://imagej.net/Fiji/Downloads). For mice cortical neurons and fly MNs ROIs of 30 μm long axon accounting for the full width of the axons were used. For human primary fibroblasts, ROI were performed marking the entire cell was used. The α-tubulin acetylation levels were extracted from mean intensity levels. Background levels were subtracted and the ratio of acetylated α-tubulin/*α*-tubulin was calculated.

### Vesicular and mitochondrial transport recording in vitro and in vivo

Axonal transport in mice cultured cortical neurons was recorded in microfluidics devices, prepared as described in (50). Briefly, air bubbles were removed from mixed sylgard 184 elastomer (VossChemie Benelux, 1:15 ratio with curing agent) by centrifuging at 1,000xg for 5 minutes. Liquid was poured into the microfluidic device mold and was cured by 3 hours incubation at 70°C incubator. Molds were cut and washed twice with 70% ethanol, air dry in biological hood and placed on 35 mm glass bottom dishes (MatTek, P35G-0-20-C). To increase the adhesiveness the microfluidic chambers and dishes were heated to 70°C, and placed on dishes. The day of the culture medium supplemented with 50 ng/ml or 20 ng/ml BDNF (PeproTech, 450-02) was added to the microfluidic devices distal or soma side, respectively. Labeling were done on after 5DIV by adding 1 μM LysoTracker® Red DND-99 (ThermoFisher Scientifics, L7528), MitoTracker® Green FM (ThermoFisher Scientifics, M7514) or MitoTracker® Deep Red FM (ThermoFisher Scientifics, M22426) 30 minutes prior to time-lapse recordings. Recording of mice cortical neurons and *Drosophila melanogaster* MNs was performed on an inverted confocal microscope Nikon, A1Ti, at 600 ms frame interval. Recordings of human primary fibroblasts were performed using in-cell 2200 (General Electric) using 60X air lens, at 2 seconds frame intervals, using temperature (37°C) and CO2 control.

Axonal transport recording in *Drosophila Melanogaster* MNs were done on 3^rd^ instar larvae expressing UAS:RNAi and Syt1-GFP. Larvae were anaesthetized with ether vapors (8 minutes) and mounted dorsally on microscope slide using 80% glycerol.

Intra-cellular transport in human primary fibroblasts were done on cultured primary fibroblasts 3 days after plating, by adding 1 μM LysoTracker® Red DND-99 (ThermoFisher Scientifics, L7528) 30 minutes prior to time-lapse recordings.

Video analysis were performed by generation of kymographs for single blind analysis using ImageJ plugin-KymoToolBox. For *Drosophila melanogaster* analysis StackReg plugin was used to align frames. Vesicles were considered stationary if speed was lower than 0.1 μm/sec.

### Plasmids and drug treatments

For silencing of *Elp3* we inserted sh-*Elp3* 5’GCACAAGGCUGGAGAUCGGUU3’ or a control sequence sh-Control 5′-TACGCGCATAAGATTAGGG-3′ previously described in (25, 51). The viral packaging vector is PSPAX2 and envelope is VSV-G. The lentiviral vector is pCDH-cmv-EF1-copGFP (CD511B1), and the promotor was replaced by a U6 promoter. For expression of human ACLY E14.5 cortical neurons in suspension were transfected with pEF6-Acly (Addgene plasmid # 70765, a gift from Kathryn Wellen) (52) or pEF6 (control) and GFP (from VPG-1001, Lonza) using Mouse Neuron Nucleofector® Kit (VPG-1001, Lonza) according to manufacturer’s protocol. GFP positive neurons were used for analysis. For expression of human ACLY cultured human primary fibroblasts were transfected with pEF6-Acly (Addgene plasmid # 70765, a gift from Kathryn Wellen) (52) or pEF6 (control) and GFP (from VPG-1001, Lonza) using Lipofectamine® 2000 according to manufacturer’s protocol. GFP positive cells were used for analysis. Tubastatin A (TBA, 20μM) or Hydroxycitrate (HCA, 3mM or 10mM) were dissolved in Ultra-Pure Water (UPW) and added to cell cultures 2 hours or 8 hours, respectively, prior to recording.

### Real Time Quantitative PCR analysis (qRT-PCR)

One mouse cortex or ten adult fly heads were collected in TRIzol Reagent (Ambion, Life Technologies) followed by RNA extraction performed using the manufacturer’s instructions. After DNAse treatment (Roche), 1 μg of RNA was reverse transcribed with RevertAid Reverse Transcriptase (Fermentas). RT-qPCR was performed using Quant Studio (Thermo) and TaqMan primers for mice, or a Light Cycler 480 (Roche) with Syber Green mix for *Drosophila melanogaster*. Analyses were done using 2-^ΔΔ^CT method (53).

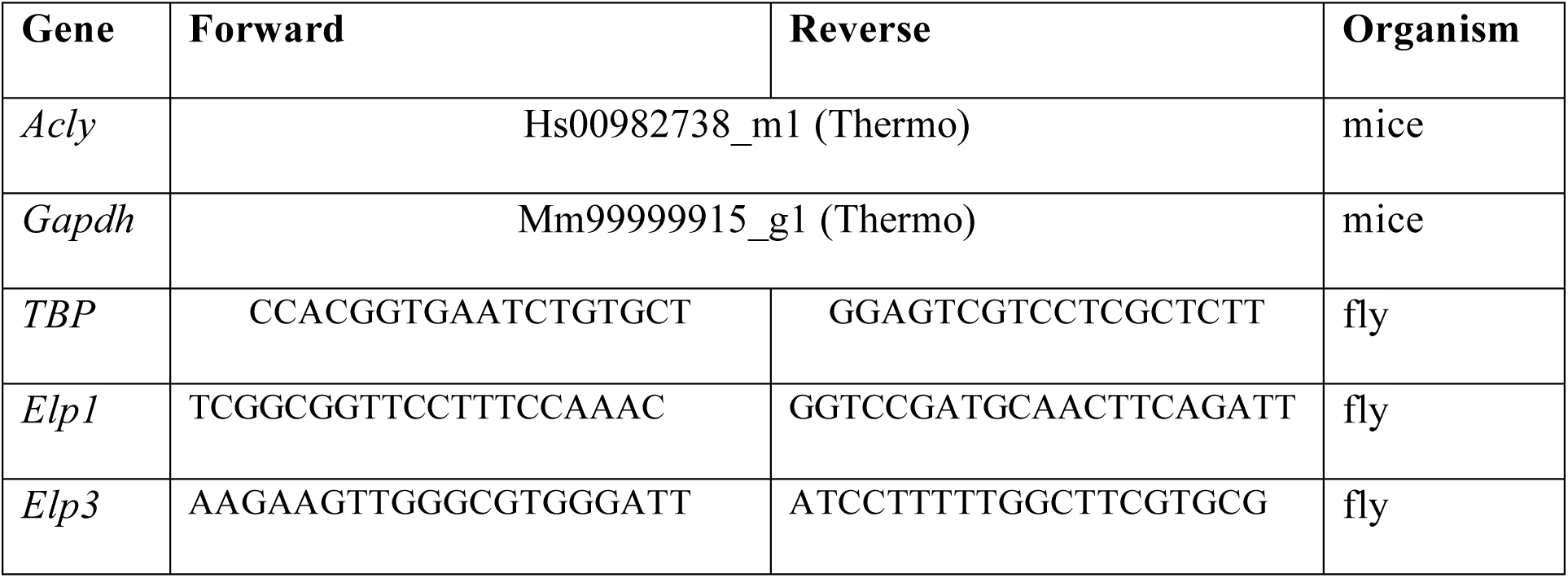

### Protein aggregation assay

Cultured mice cortical neurons or human primary fibroblasts at 60% confluency were incubated with PROTEOSTAT® (Enzo LifeSciences) to detect protein aggregates.

### Locomotion activity and climbing assays

Larval crawling speed assays were performed by placing 3^rd^ instar larvae in the center of 15-cm petri dishes coated with 3% agar as previously published (54), velocities were extrapolated by measuring the distance traveled in one minute. Climbing assay were performed as previously described in (55), by measuring the average ratio of successful climbs over 15 cm for 10 adult flies.

### Western blot

Mouse brain cortices, adult fly heads or human primary fibroblasts were quickly homogenized on ice in RIPA buffer, or 320 mM sucrose, 4 mM HEPES buffer for subcellular fractions. Protease inhibitor cocktail (Roche, P8340 or Sigma-Aldrich, S8820) and 5 μM Trichostatin-A (Sigma-Aldrich, T8552) to inhibit protein degradation and *α-*tubulin deacetylation. 2 μg of protein lysate were used for α-tubulin acetylation analysis and 20-30 μg for all other proteins. Nitrocellulose membranes were imaged using Amersham Imager 600 (General Electric, 29083461) and band densitometry was measured using FIJI.

| **Protein** | **Company** | **Cat #** | **WB** | **IF** | **ELISA** |
| --- | --- | --- | --- | --- | --- |
| α-Tubulin | Sigma-Aldrich | T9026 | 1:5,000 | | 1:2,000 |
| $\alpha/\beta$-Tubulin<br>(total tubulin) | Cytoskeleton | ATN02-A | | 1:150 | |
| acetylated α-Tubulin | Sigma-Aldrich | T7451 | 1:15,000 | 1:15,000 | |
| β-Tubulin | cell signaling | 2146 |  | 1:100 |  |
| β-actin | Sigma-Aldrich | A3853 | 1:20,000 |  |  |
| HDAC6 | Santa Cruz<br>Biotechnology | sc-5258 | 1:200 |  |  |
| ATAT1 | Max Nachury |  | 1:1000 |  |  |
| Tau-1 | Millipore | MAB3420 |  | 1:500 |  |
| CSP | DSHB | DCSP-2<br>(D6D) |  | 1:10 |  |
| ELP1/IKAP | Anaspec | AS-54494 | 1:500 |  |  |
| ELP3 | Jesper Svejstrup |  | 1:1,000 |  |  |
| Synaptophysin | Sigma-Aldrich | s5768 | 1:1,000 |  |  |
| ACLY | cell signaling | <u>13390</u> | 1:1,000 |  |  |
| goat anti mouse | Jackson<br>ImmunoResearch<br>Labs | 115-035-003 | 1:10,000 | 1:200 | 1:5,000 |
| goat anti rabbit | Jackson<br>ImmunoResearch<br>Labs | 111-035-003 | 1:10,000 | 1:200 |  |
| donkey anti goat | Jackson<br>ImmunoResearch<br>Labs | 705-035-003 | 1:10,000 |  |  |

### Subcellular Fractionation

Subcellular fractionation of frozen mice brain cortex was performed as previously described (30).

### Mass Spectrometry Analysis

Pellets from three independent samples of pooled vesicles isolated from brain cortex of WT or *Elp1* cKO mice were solubilized using 5% SDS. The samples were dissolved in 10mM DTT 100mM Tris and 5% SDS, sonicated and boiled in 95^°^ C for 5 minutes. The samples were precipitated in 80% acetone. The protein pellets were dissolved in in 9M Urea and 100mM ammonium bicarbonate than reduced with 3mM DTT (60°C for 30 min), modified with 10mM iodoacetamide in 100mM ammonium bicarbonate (room temperature for 30 minutes in the dark) and digested in 2M Urea, 25mM ammonium bicarbonate with modified trypsin (Promega), overnight at 37°C in a 1:50 (M/M) enzyme-to-substrate ratio. The resulting tryptic peptides were desalted using C18 tips (Harvard) dried and re-suspended in 0.1% Formic acid. They were analyzed by LC-MS/MS using Q Exactive plus mass spectrometer (Thermo) fitted with a capillary HPLC (easy nLC 1000, Thermo). The peptides were loaded onto a homemade capillary column (20 cm, 75 micron ID) packed with Reprosil C18-Aqua (Dr Maisch GmbH, Germany) in solvent A (0.1% formic acid in water). The peptides mixture was resolved with a (5 to 28%) linear gradient of solvent B (95% acetonitrile with 0.1% formic acid) for 60 minutes followed by gradient of 15 minutes gradient of 28 to 95% and 15 minutes at 95% acetonitrile with 0.1% formic acid in water at flow rates of 0.15 μl/min. Mass spectrometry was performed in a positive mode using repetitively full MS scan followed by high collision induces dissociation (HCD, at 35 normalized collision energy) of the 10 most dominant ions (>1 charges) selected from the first MS scan. The mass spectrometry data was analyzed using the MaxQuant software 1.5.1.2. (www.maxquant.org) using the Andromeda search engine, searching against the mouse uniprot database with mass tolerance of 20 ppm for the precursor masses and 20 ppm for the fragment ions. Peptide- and protein-level false discovery rates (FDRs) were filtered to 1% using the target-decoy strategy. Protein table were filtered to eliminate the identifications from the reverse database, and common contaminants and single peptide identifications. The data was quantified by label free analysis using the same software, based on extracted ion currents (XICs) of peptides enabling quantitation from each LC/MS run for each peptide identified in any of experiments.

### In vitro α-tubulin assay

*In vitro* α-tubulin acetylation assay was performed as previously described (30).

### α-Tubulin deacetylation assay

α-Tubulin deacetylation assay was performed as previously described in (38). 96 well half area plates (Greiner, 674061) were coated with 1 μg tubulin in 50 μl of ultra-pure water for 2.5 hours at 37°C, followed by blocking (PBS, 3% BSA, 3% skim milk, 3% fetal bovine serum) for 1 hour at 37°C, and washing with PBS+0.05% Tween-20. 10 μg of cytosolic fraction isolated from newborn mice brain cortex in deacetylation buffer (50 mM Tris-HCl at pH 7.6, 120 mM NaCl, 0.5 mM EDTA) with protease inhibitor cocktail and phosphatase inhibitor (to avoid dephosphorylation of HDAC6), were added per well for incubation of 4 hours at 37°C with shaking at 100 RPM. The wells were washed and incubated overnight at 4°C with acetylated α-tubulin antibody (1:2,000) in blocking buffer (PBS+0.05% Tween-20 +3%BSA), wells were washed and incubated for 2 hours at 37°C with Peroxidase conjugated Goat anti mouse antibody (1: 5,000) in antibody blocking buffer, following another wash samples were incubated with TMB/E (ES001, Merck Millipore), reaction was stopped with H_2_SO_4_.

### ACLY activity assay

ACLY activity assay was measured as previously described in (41). Cells or brain extracts were disrupted by passing them 15 times in 25-gauge needle in 100 mM Tris-HCL buffer, supplemented with protease and phosphatase inhibitors. 5ug of cells/brain lysate were added to reaction mix (200 mM Tris-HCL PH 8.4, 20 mM MgCl2, 20 mM sodium citrate, 1 mM DTT, 0.1 mM NADH (Sigma-Aldrich, N8129), 6 U/mL Malate dehydrogenase (Sigma-Aldrich, M1567), 0.5 mM CoA (Sigma-Aldrich, C3019) with or without ATP (Sigma-Aldrich, A1852). ACLY activity was measured every 100 seconds for 4 hours, in volume of 50ul in 384 well plate using 340nm OD read. ACLY specific activity was calculated as the change in absorbance with ATP compared to the change without ATP. For statistical comparison ACLY activity was define as the slope from the linear range of the reaction.

### Acetyl-CoA sample preparation and LC–MS/MS analysis

Acetyl-CoA was extracted as previously described (56). Briefly, cold methanol (500 μl; -20 °C) was added to the cell pellets, and the mixture was shaken for 30 s (10 °C, 2000 r.p.m., Thermomixer C, Eppendorf). Cold chloroform (500 μl; −20 °C) was added, the mixture was shaken for another 30 seconds, and then 200 μl of water (4 °C) was added. After the mixture was shaken for 30 seconds and left on ice for 10 minutes, it was centrifuged (21,000×g, 4 °C, 10 min). The upper layer was collected and evaporated. The dry residue was re-dissolved in eluent buffer (500 μl) and centrifuged (21,000×g, 4 °C, 10 minutes) before placing in LC-MS vials. Acetyl CoA was analyzed as previously described (57). Briefly, the LC–MS/MS instrument consisted of an Acquity I-class UPLC system and Xevo TQ-S triple quadrupole mass spectrometer (both Waters) equipped with an electrospray ion source. LC was performed using a 100 × 2.1-mm i.d., 1.7-μm UPLC Kinetex XB-C18 column (Phenomenex) with mobile phases A (10 mM ammonium acetate and 5 mM ammonium hydrocarbonate buffer, pH 7.0, adjusted with 10% acetic acid) and B (acetonitrile) at a flow rate of 0.3 ml min-1 and column temperature of 25°C. The gradient was as follows: 0-5.5 minutes, linear increase 0–25% B, then 5.5-6.0 minutes, linear increase till 100% B, 6.0-7.0 minutes, hold at 100% B, 7.0-7.5 minutes, back to 0% B, and equilibration at 0% B for 2.5 minutes. Samples kept at 4°C were automatically injected in a volume of 5 μl. Mass spectrometry was performed in positive ion mode, monitoring the MS/MS transitions m/z 810.02 → 428.04 and 810.02 → 303.13 for acetyl-CoA. Spikes of defined amounts of AcCoA were added to the samples to confirm the absence of signal inhibition (matrix effect) in the analyzed extracts. Quantification of AcCoA was done against external calibration curve with 1–1,000 ng ml–1 range of AcCoA concentrations using TargetLynx software (Waters).

### Analysis and statistics

All experiments were performed under single blinded condition and statistical analyses were generated with GraphPad Prism Software 7.0

## Author contributions

A.E., G.M., M.W., and L.N. designed the study. A.E. and G.M. performed and interpreted most experiments. R.L.B. contributed to *Drosophila* work. M.S. contributed to cell cultures and biochemical work. S.T. and L.B. performed biochemical experiments with Elp3cKO embryos. A.B. performed acetyl co-A LC-MS/MS analysis. Shani Inbar performed qPCR analysis. P.D. and I.D. maintained Elp1 cKO mice colonies and provided brain material. F.S., A.C., B.B, and J.M.R provided guidance and help for experiments. M.W. and L.N. contributed to data interpretation; and A.E., G.M., M.W., and L.N. wrote the manuscript with input from all coauthors.

## Competing interests

The authors declare that they have no competing interests.

## Acknowledgements

We thank Maria M. Magiera and Carsten Janke for providing non acetylated Hela Tubulin. We thank Patrik Verstreken for sharing Elp3KD flies, M. Nachury for sharing ATAT1 antibody and K. Sadoul for providing the Atat1 KO mice, T. Lahusen from American Gene Technologies for creating viral sh-RNA particles and E. Even for graphical design. We are grateful to Francesca Bartolini and Marina Mikhaylova for their constructive feedback on the manuscript as well as to all members of the Nguyen and Weil laboratories for their critical reading. L.N. is Senior Research Associates from F.R.S-F.N.R.S. This work was supported by the F.R.S.-F.N.R.S. (Synet; EOS 0019118F-RG36), the Fonds Leon Fredericq (L.N.), the Fondation Médicale Reine Elisabeth (L.N.), the Fondation Simone et Pierre Clerdent (L.N), the Belgian Science Policy (IAP-VII network P7/20 (L.N.)), and the ERANET Neuron STEM-MCD and NeuroTalk (L.N.); grants from Agence Nationale de la Recherche (ANR-18-CE16-0009-01 AXYON (F.S.); ANR-15-IDEX-02 NeuroCoG (F.S.) in the framework of the “Investissements d’avenir” program); Fondation pour la Recherche Médicale (FRM, DEI20151234418, F.S.). A.E.’s stay at GIGA Research Institute of the University of Liège was funded by EMBO Short-Term Fellowships (ASTF 174-2016), A.E., M.S., and M.W.’s research was supported by the Israel Science Foundation (grant no. 1688/16). The authors declare no competing financial interests.

## Notes

### Competing Interest Statement

The authors have declared no competing interest.

